# Shapley Fields Reveal Chemotopic Organization in the Mouse Olfactory Bulb Across Diverse Chemical Feature Sets

**DOI:** 10.1101/2025.02.26.640432

**Authors:** Nikola Milićević, Shawn D. Burton, Matt Wachowiak, Vladimir Itskov

## Abstract

Representations of chemical features in the neural activity of the olfactory bulb (OB) are not well-understood, unlike the neural code for stimuli of the other sensory modalities. This is because the space of olfactory stimuli lacks a natural coordinate system, and this significantly complicates characterizing neural receptive fields (tuning curves), analogous to those in the other sensory modalities. The degree to which olfactory tuning is spatially organized across the OB, often referred to as chemotopy, is also not well-understood. To advance our understanding of these aspects of olfactory coding, we introduce an interpretable method of Shapley fields, as an olfactory analog of retinotopic receptive fields. Shapley fields are spatial distributions of chemical feature importance for the tuning of OB glomeruli. We used this tool to investigate chemotopy in the OB with diverse sets of chemical features using widefield epifluorescence recordings of the mouse dorsal OB in response to stimuli across a wide range of the chemical space.

We found that Shapley fields reveal a weak chemotopic organization of the chemical feature sensitivity of dorsal OB glomeruli. This organization was consistent across animals and mostly agreed across very different chemical feature sets: (i) the expert-curated PubChem database features and (ii) features derived from a Graph Neural Network trained on human olfactory perceptual tasks. Moreover, we found that the principal components of the Shapley fields often corresponded to single commonly accepted chemical classification groups, that therefore could be “recovered” from the neural activity in the mouse OB. Our findings suggest that Shapley fields may serve as a chemical feature-agnostic method for investigating olfactory perception.

## Introduction

Primary sensory areas of the mammalian brain, such as the visual and the auditory systems, are known to encode characteristic features of the stimuli, such as the wavelength of light and frequency of sound. Furthermore, encoding of such stimulus features is spatially organized within topographic maps across primary and downstream cortical areas of visual, auditory, gustatory, and other sensory systems Chen et al. (2011); Hubel and Wiesel (1977); Kandler et al. (2009). In contrast to these sensory systems, however, how physicochemical features of olfactory stimuli are encoded within the olfactory system, including whether they are topographically organized within the olfactory bulb (OB) – the primary olfactory sensory area in the mammalian brain – remains poorly understood.

Axons of the olfactory sensory neurons (OSNs) expressing the same olfactory receptor (OR) gene converge to deliver sensory input from the periphery to stereotypically positioned glomeruli on the OB surface. Due to the large number of OR genes, the dimensionality of olfactory stimulus space is potentially quite large. However, dimensionality reduction has been found to account for odorant similarity in chemical, neuronal and perceptual spaces in humans Koulakov (2011); Keller and Vosshall (2016), mice Ma et al. (2012) and across species Haddad et al. (2008, 2010); Saito et al. (2009). These results indicate that olfaction follows a combinatorial logic rather than a simple labeled-line code, similar to receptive field patterns in lower-dimensional sensory systems.

Several studies indicate that glomeruli in mammalian OBs tuned to similar combinations of odorant chemical features are also located in close proximity on the OB and form chemical-feature clusters Burton et al. (2022); Mori et al. (2006). For example, in the rat OB, a homologous series of odorant molecules can evoke distinct but spatially clustered responses that change systematically with carbon chain length Johnson et al. (1999); Johnson and Leon (2000); Meister and Bonhoeffer (2001); Rubin and Katz (1999). Moreover, glomerular responses to certain odorant functional groups and hydrocarbon substructures also follow a systematic organization in the rodent OB Johnson et al. (2005); Johnson and Leon (2007); Matsumoto et al. (2010); Soelter et al. (2020); Takahashi et al. (2004); Uchida et al. (2000). These findings provide evidence for a topographic mapping of chemical features – or “chemotopy” – in glomeruli of the OB.

Several studies have cast doubt on the validity of the chemotopic organization, however. It was found in Ma et al. (2012); Chae et al. (2019); Soucy et al. (2009) that commonly used physicochemical features were not systematically represented by glomerular activity in the mouse OB. Instead, Ma et al. (2012) argued that there is no chemotopic representation of chemical features in either fine or broad scales, while there is still a significant correlation between pairwise tuning similarity and distances between glomeruli. In Burton et al. (2022); Chae et al. (2019); Soucy et al. (2009) the authors argued for a functional “mosaic” organization of glomerular activity.

It is natural to define weak chemotopy as the property by which glomeruli, tuned to chemical features in “close proximity” in chemical feature space, are also in a spatial proximity on the OB. We call this chemotopy “weak” in reference to the imprecise and overlapping nature of the topographic organization with respect to the chemical features. This is in contrast to retinotopy in the primary visual cortex, where the visual receptive fields are sharply localized. However, the nature of chemotopy in the OB seems to greatly depend on the choice of chemical features, because different choices of chemical features significantly change the distances in the poorly understood chemical space. This dependence on the considered features may explain the inconsistencies of the previous studies of chemotopy.

Here, we set out to characterize chemotopy and its relationship to the choice of the chemical features that are being represented in the neural responses. To this end we developed a novel method of Shapley fields, as an olfactory analog of the retinotopic receptive fields. Shapley fields are distributions of chemical feature importance for the tuning of OB glomeruli. They are derived as an aggregate from Shapley values, a machine learning tool that allocates credit to features that explain model predictions Lundberg and Lee (2017), originally introduced in the context of game theory Shapley (1951). Another reason to use Shapley fields is that this tool automates feature selection and thus allows analysis of olfactory receptive properties using any significantly diverse chemical feature set. We used this tool to characterize chemotopy relative to diverse sets of chemical features using epifluorescence recordings of the mouse dorsal OB in response to stimuli across a wide range of odorants.

Our findings are consistent with a weak chemotopic organization of odor representations across OB glomeruli. We also established that our analyses do not depend on different choices of chemical feature sets: (i) the expert-curated PubChem chemical feature set and (ii) an artificial chemical feature set produced by a neural network, trained on human perception and molecular graphs. Both feature sets captured similar receptive properties in the mouse OB.

## Results

Odorant-evoked responses were imaged with widefield epifluorescence across the dorsal surface of the OB in anesthetized mice, using artificial inhalation to ensure consistent odorant sampling (Figure 1A), see Methods and Burton et al. (2022).

**Fig. 1:**
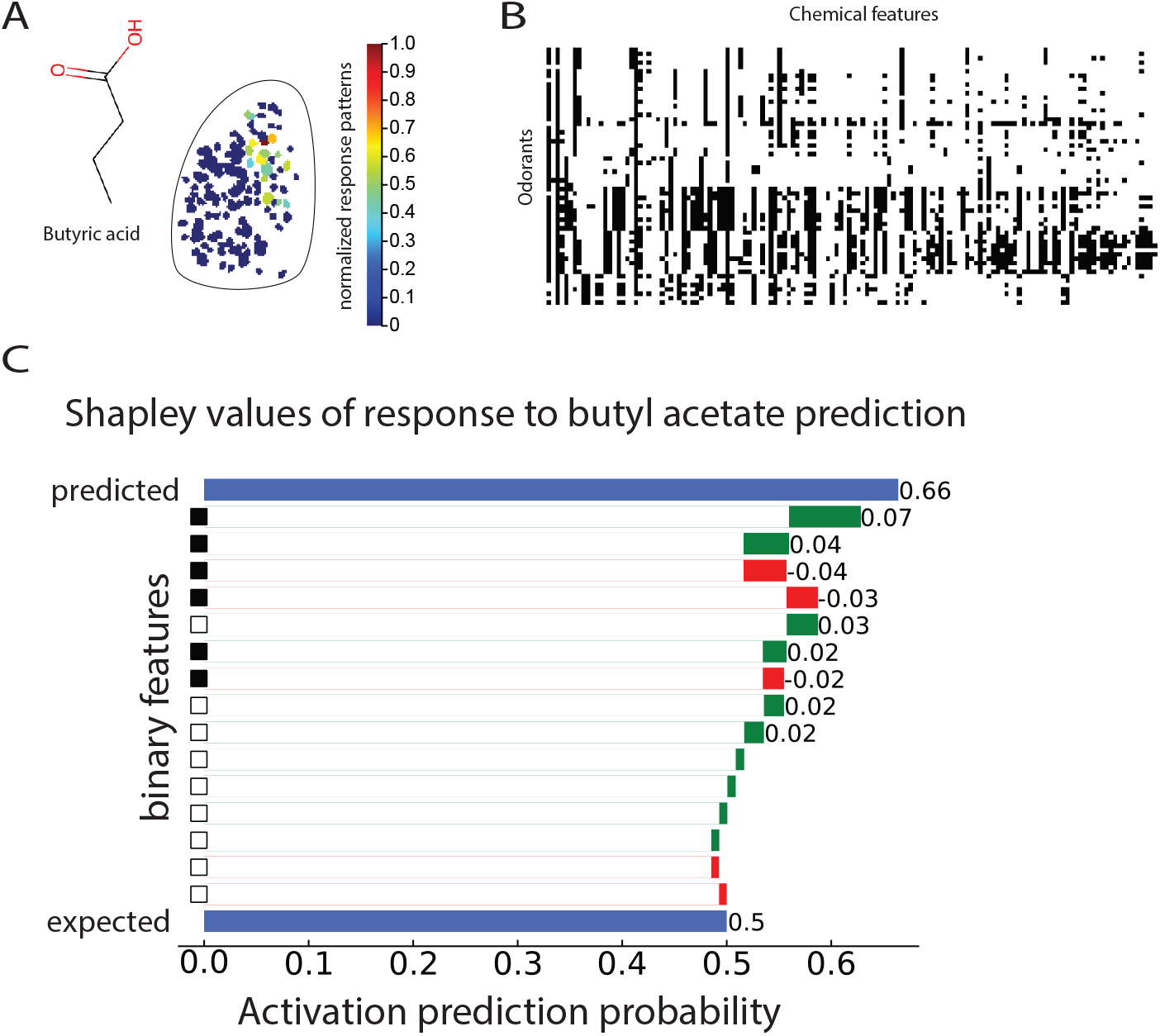
Predicting glomerular activation in response to odorants. **A. Glomerular activation**. An example of glomerular activation pattern in response to butyric acid. **B. PubChem chemical features**. Binary chemical features of the presented odorants, from the PubChem database (Methods). Black indicates presence of a chemical feature, while white indicates absence. **C. Explanation of chemical feature contribution towards predicting activation of a glomerulus to a particular odorant (butyl acetate)**. The contribution of each feature towards model prediction is calculated with Shapley values (Methods). Positive Shapley values (green) show contribution towards predicting activation; negative Shapley values (red) show contribution towards predicting no activation. The black squares on the *y*-axis indicate presence of a particular chemical feature in the odorant, the white squares indicate absence. The bottom blue line is the expected prediction; probability of activation the model would predict if no chemical features were given to the model. In this example, all Shapley values of features together with the model expected prediction sum to give the 66% probability of activation (top blue bar). This is represented in the waterfall plot; each feature’s Shapley value contributes to the model prediction starting from 0.5 and moving the prediction to the left (if Shapley value is negative) or the right (if Shapley value is positive). Only the 15 features with the largest absolute Shapley values are shown in decreasing order from top to bottom.

To evaluate the sensitivity of glomeruli to odorant chemical features, we trained a Chemotuning model using the PubChem binary molecular fingerprint features (Figure 1B), Kim et al. (2022). The Chemotuning model assumes that each glomerulus activates based on a set of chemical features of a given odorant stimulus, irrespective of the glomerulus position in the OB. For each glomerulus, a random forest binary classifier was fit to predict activation of the glomerulus in response to the odorant, from the chemical features. More precisely, we trained a classifier to only detect activation or no activation. Our analysis did not consider whether a particular odorant activated a glomerulus more than another odorant (see Methods).

To understand the chemical receptive properties of glomeruli, we used Shapley values. Shapley values are a machine learning tool that enable allocation of credit to features explaining model predictions Lundberg and Lee (2017). This tool is inspired by classical work in game theory Shapley (1951); our usage is similar to that in Zoabi et al. (2021), and Rodríguez-Pérez and Bajorath (2019). Shapley values are a general framework that unifies several other popular additive feature attribution methods such as LIME Ribeiro et al. (2016), DeepLIFT Shrikumar et al. (2017) and Layer-wise relevance propagation Bach et al. (2015). We used mean absolute Shapley values, similar to Lundberg and Lee (2017) (see Methods), to determine which chemical features are most responsible for predicting glomerular activation by any given odorant (Figure 1C).

### Shapley fields reveal a stereotypical mosaic organization of chemical receptive fields within the olfactory bulbs

To understand the organization of OB chemical tuning, we constructed Shapley fields. Shapley fields represent signed distributions of feature credit allocations for prediction of glomerular activation, mapped on the surface of the OB. In other words, they are a signed projection of odorant chemical feature importance according to the Chemotuning model on the OB. Alternatively, a Shapley field of a chemical feature is the vector of Shapley values for each glomerulus. Shapley fields associated with four of the PubChem features are shown in Figure 2A.

**Fig. 2:**
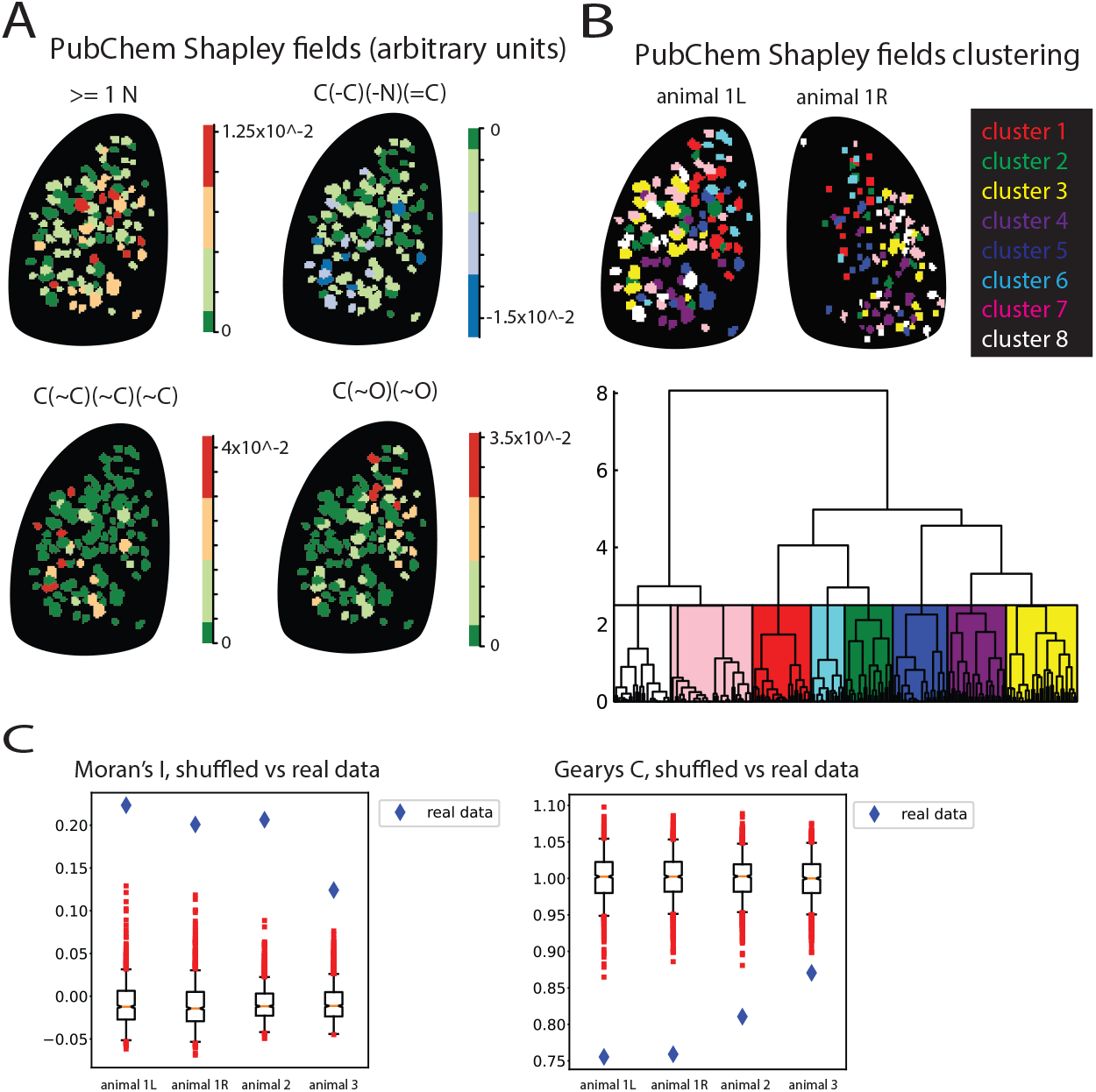
Mosaic chemotopic organization of feature importance on the OB. **A. Shapley fields**. Importance of four chemical features from the PubChem feature set is displayed on the surface of the left OB, of animal 1, via Shapley fields (Methods). The top right figure is the negative Shapley field of the feature C(-C)(-N)(=C). The rest of the figures are the positive Shapley fields of the respective features. **B. Clustering of Shapley fields results in spatially correlated clusters on the surface of OB**. Hierarchical clustering of glomeruli on left and right OBs of animal 1, based on the first 20 principal components of Shapley fields, for the Pubchem chemical features. Cosine distance ward linkage clustering was used on the principal components of Shapley fields. Most clusters have a “mosaic” distribution that is roughly consistent between the left and the right OBs. **C. The spatial clustering inferred from Shapley fields is statistically significant**. Glomeruli positions on the OB were randomly shuffled n times (*n* = 1000). After each shuffling, a glomerulus was assigned to its original cluster from Panel B. The Moran’s I and Gearys C clustering scores (Methods), across all 4 datasets show a highly significant difference (p=0) between the clusters from the data (blue) and the randomly shuffled ones (red).

To investigate the spatial organization of the feature importance on the surface of the OB, we computed the 20-dimensional PCA projections of Shapley fields (see Methods) and then performed a hierarchical clustering of glomeruli, based on the values of Shapley fields after the projection assigned to the glomeruli (Figure 2B). The resulting clusters revealed a mosaic organization of glomeruli across the left and the right OBs of an animal. This organization was inferred from the Shapley values of our Chemotuning model. This organization was also observed across animals and with respect to a different chemical feature set (SuppFig 1A-C).

Notably, while the Chemotuning model does not consider glomerular positional data, spatial clustering by chemical feature responsivity nevertheless emerged. This is similar to the clustering previously reported by some studies Burton et al. (2022); Johnson and Leon (2007); Mori et al. (2006). This organization proved highly statistically significant (Figure 2C), as revealed by shuffling the identity of glomeruli (*N* = 1000) and computing spatial-autocorrelation statistics, Moran’s I and Geary’s C (see Methods) of the original clustering and the clustering after each shuffling. The same was true for a different set of chemical features (SuppFig 1D). For the sake of clarity, we displayed the analysis using 8 clusters, however any choice of number of clusters between 6 and 10 was statistically significant (*p <* 0.05) with respect to autocorrelation scores of Moran’s I and Geary’s C (SuppFig 2).

### The Chemotopy model is more predictive than the Chemotuning model

The Chemotuning model prediction does not take advantage of the position of glomeruli on the OB. If there is a spatial organization based on chemistry, then adding positional information as a feature for learning should improve the model’s performance. To this end, we also developed the Chemotopy model. The Chemotopy model assumes that the response of a glomerulus is a function of both position and the chemical features of the odorant and predicts glomerular activation from both the chemical features of the odorant as well as the positional information of the glomerulus. The Chemotopy model is thus a single random forest model for predicting activation of all the observed glomeruli on the OB.

We compared the predictions of the Chemotuning and the Chemotopy models, as well as two different baseline models. Here we considered the baseline of predicting that the glomerulus would not be activated (because glomerular activation is sparse in the dataset) as well as the peer prediction model (see below). In order to have a relatively small number of model parameters, we used the principal components of the PubChem features (“PCA-PubChem”), while ordering them according to their Shapley values (see Shapley rankings in Methods). The Chemotopy model outperformed the Chemotuning model when we used the Shapley ranking of the PCA-PubChem features. The Shapley rankings are stable with respect to the randomness inherent in our random forest models, see Methods and SuppFig 3. To provide more context to this comparison, we additionally considered a peer prediction model as the other baseline.

Our peer prediction model (see Methods) is inspired by Harris et al. (2003); Itskov et al. (2008); McKenzie et al. (2021); Luczak et al. (2022),where it has been observed that one could predict spiking of individual neurons based on the activity of other neurons. In our data, the peer prediction model provided high prediction accuracy (Figure 3) because the olfactory representations are likely redundant for the stimuli that we considered, as should be expected from combinatorial coding of olfactory stimuli in the OB Malnic et al. (1999); Koulakov et al. (2006); de March et al. (2020); Grabe and Sachse (2018). We observed that approximately 70 − 80% of the difference between the peer prediction model’s quality of prediction and the baseline is accounted for by the chemical feature sets that we used. Of note, our odorant sample size is rather small (*n* = 59) as compared to the number of recorded glomeruli in a single OB (*m* = 113, 108, 128, 132). Thus, we expect the quality of prediction of Chemotopy and Chemotuning models to catch up to the peer prediction model’s performance with a significantly larger sample of odorants. Note that the analysis in Figure 3 was for the response data on the left OB of animal 1, however the same observations held for all the other OBs in our data (SuppFig 4).

**Fig. 3.**
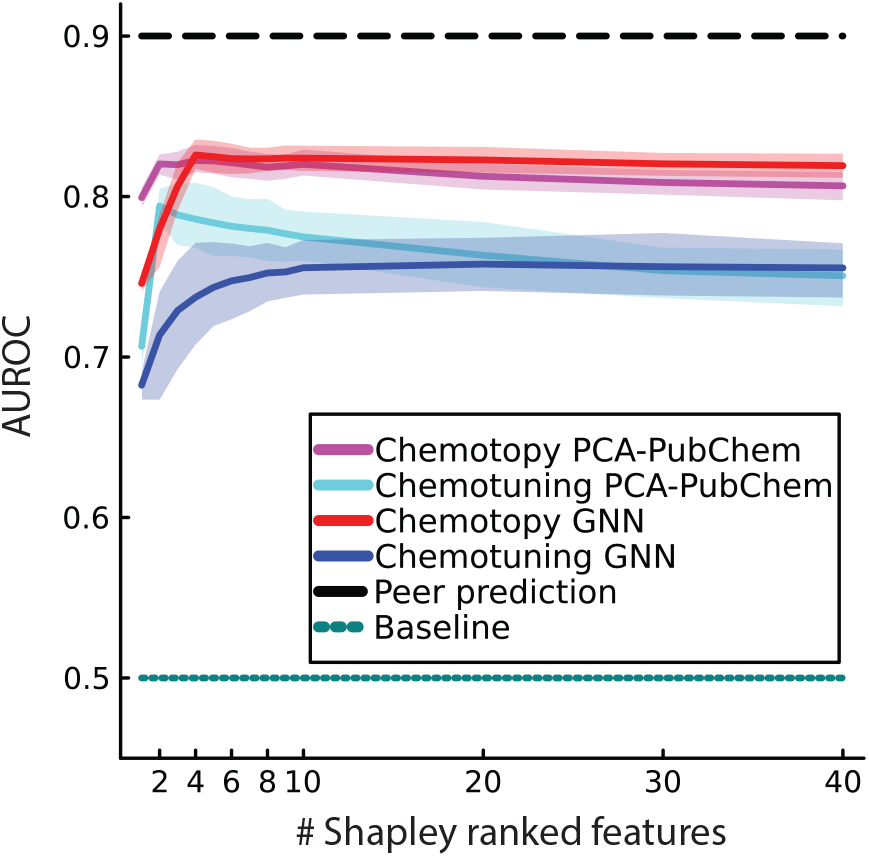
Chemotopy model outperforms chemotuning model. Three different models: chemotuning, chemotopy and the peer prediction are compared vs baseline, across chemical feature sets. The features in a chemical feature set were ranked according to their global Shapley values, largest global Shapley value first (Methods). The features were added, one at a time according to their Shapley ranks, to the two models of glomerular activation prediction from chemical features; chemotopy and chemotuning (Methods). The mean cross-validated AUROC score is plotted along with its 95% confidence interval (shaded regions), *N* = 200, against the number of Shapley ranked chemical features used. The mean cross-validated peer prediction model (black dashed line) AUROC score is also shown as well as the baseline (teal) having value of 0.5 which is the AUROC score of a random binary classifier. The 95% confidence interval for the peer prediction was too narrow to be perceptible on the plot (0.8970 *−* 0.9026).

Given the previously discussed discrepancies in reported results, which may arise from the differing chemical feature sets used, we sought to explore whether the superior performance of the Chemotopy versus Chemotuning models was influenced by the specific choice of chemical features. To investigate this, we also did our analysis using a second set of chemical features. We generated these features from a graph neural network (GNN) trained on human derived odor labels of approximately 6000 odorant molecules from the GoodScents company Company (2023), similar to how Lee et al. (2023); Achebouche et al. (2022); Qian et al. (2023) learned odorant representations. We used the same GNN architecture as was reported in Achebouche et al. (2022) (see Methods). The Chemotopy model significantly outperformed the Chemotuning model, with the accuracy gap widening as more Shapley-ranked features were included (Figure 3). Notably, the Chemotopy model’s prediction accuracy peaked at just five features, whereas the Chemotuning model showed no clear peak. Here, the Shapley values served as a dimensionality reduction technique that improved the quality of prediction by only considering the most relevant features.

We also found that GNN features do not outperform the PCA-PubChem feature set with respect to predicting glomerular activation (Figure 3). This finding contrasts with recent works Achebouche et al. (2022); Qian et al. (2023) where GNN features have been reported to outperform other feature sets such as RDKit Landrum (2010) across different olfaction datasets and olfaction tasks (across species). Comparing models trained on GNN features with models trained on PCA-PubChem, we see that PCA-PubChem has more predictive power than the GNN features early on. However, as more features get added, GNN features match PCA-PubChem predictive power both in the Chemotopy and Chemotuning models (Figure 3).

### Chemical features derived from human perception yield predictions and mosaic chemotopic organization that is similar to those derived from the PubChem chemical features

How do the chemical receptive properties of the glomeruli depend on the choice of the chemical feature sets? To answer this question, we first computed the Shapley fields of GNN features (Figure 4A). The first principal component of the GNN Shapley fields was highly correlated to the glomerular carboxylic acid sensitivity vector, which counts how many acids each glomerulus responded to (between 0 and 5 in the stimulus set). Tuning to acids was also correlated to several principal components of the PubChem Shapley fields, albeit to a lesser extent (Figure 4B). Note that these are the Shapley fields of PubChem features, and not PCA-PubChem features. This was also true for the OBs where the response to amines was discarded from the analysis (Methods, SuppFig 5A,B).

**Fig. 4:**
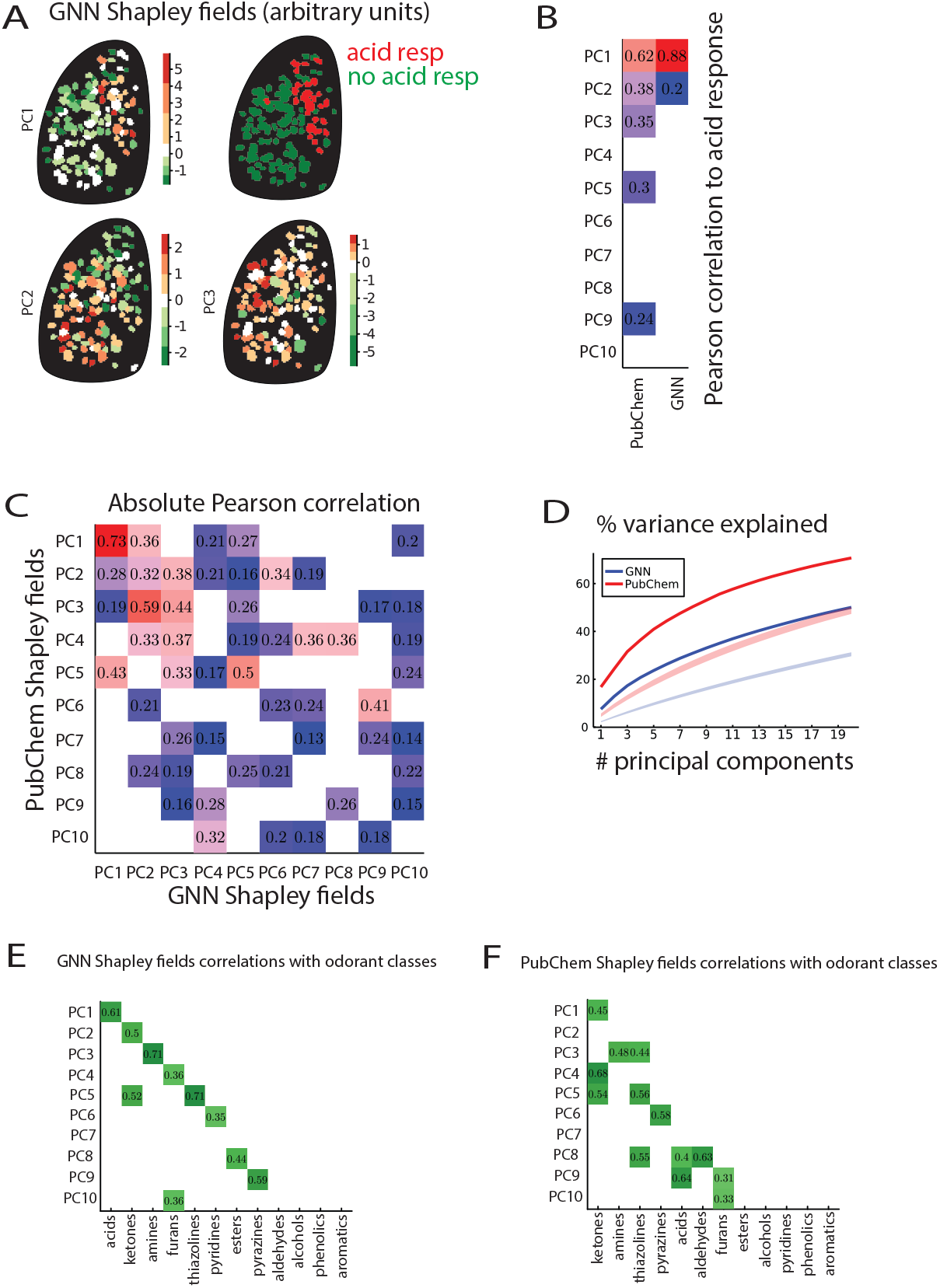
Shapley fields reveal an underlying “universal” chemical organization on the OB. **A. Principal components of Shapley fields**. Importance of GNN features is projected on the OB via Shapley fields (Methods). First 3 principal components of Shapley fields on the left OB of animal 1, along with a classification of glomeruli into acid responsive (red) and acid non-responsive (green) are displayed. Visually the first principal component of GNN Shapley fields corresponds to acid tuning separation on the OB. **B. Interpretation of principal components of Shapley fields**. Absolute Pearson correlation values between first 10 principal components of PubChem and GNN Shapley fields with the acid sensitivity vector. The acid sensitivity vector is a count of the number of acids that activate a given glomerulus. Only significant correlations (*p <* 0.05) are shown. GNN Shapley fields PC1 is highly correlated with responsiveness to acids (*r* = 0.88). All the principal components of the GNN Shapley fields beyond PC2 show no significant correlation with acid sensitivity vector. In contrast, several PubChem Shapley fields principal components show correlation to the acid sensitivity vector to some degree, albeit none as high as the PC1 of GNN Shapley fields. **C. The Shapley fields of PubChem features and GNN features are highly correlated**. The absolute values of Pearson correlations of GNN’s and PubChem’s first 10 principal components of the respective Shapley fields are displayed. Only significant correlations (*p <* 0.05) are shown. There are significant degrees of correlation among the 10 principal components of Pubchem and GNN Shapley fields. **D. Variance explained by principal components**. We find that the first 20 principal components of PubChem Shapley fields account for about 70% of the total variance in Shapley fields (red curve). In contrast, the first 20 principal components of GNN Shapley fields account for about 50% (blue curve). The most probable reason for this difference is the simple difference in the number of Shapley fields; there are 280 PubChem Shapley fields and 512 GNN Shapley fields. The percentage of variance explained is statistically significant when compared to the shuffled Shapley fields principal components, from panel D, for both PubChem (red shaded region) and GNN (blue shaded region). **E. Principal components of Shapley fields are correlated with sensitivities of** *≤* **2 odorant classes**. Principal components of GNN Shapley fields were projected onto the linear subspace spanned by odorant classes sensitivity vectors on the OB (Methods). The projection of principal components of Shapley fields can be written as a linear combination of different odorant classes sensitivity vectors. Statistical analysis involving the shuffling of odorant identities (*N* = 10000) determined which of the coefficients in the projection are statistically significant (*p <* 0.05), see Methods. Only the absolute values of statistically significant coefficients, for each projection, are shown in the table. **F. Principal components of PubChem fields are correlated with sensitivities of different odorant classes on the OB**. Same analysis as in panel E, but for the PubChem feature set.

Pearson correlation analysis also revealed significant correlations among the first ten principal components of both PubChem and GNN Shapley fields (Figure 4C, SuppFig 5C). To validate the apparent correlations between multiple principal components of GNN and PubChem Shapley fields in Figure 4C, we computed the chordal distances between subspaces generated by up to *k* = 20 principal components of each chemical feature set (see Methods). The chordal distances between the subspaces of different chemical features are significantly lower than expected from random controls (SuppFig 6). Moreover, we found that 20 principal components of the Shapley fields captured 70% of variance for the PubChem feature set and 50% of variance for the GNN feature set respectively (Figure 4D, SuppFig 5D). This suggests that the Shapley fields represent approximately the same chemical sensitivity irrespective of very different choices of the chemical features that we used: (i) a collection of the chemical properties, and (ii) features derived from human perception and molecular graphs. It is thus likely that Shapley fields capture roughly the same information about receptive properties of the OB, from any diverse enough set of chemical features.

### Principal components of Shapley fields recover major chemical classes

The *n* = 59 odorants that we used can be organized into 12 chemical classes (+1 that is the “mixed” class), as defined by major substructural motifs (see SuppFig 7 for the class assignments). To understand how the Shapley fields relate to this common chemical classification of odorants, we extended our earlier analysis (Figure 4A,B) to examine how the principal components of Shapley fields correlated with glomerular responses to all other odorant classes, by considering the odorant class sensitivity vectors. An odorant class sensitivity vector represents the sensitivity of all the recorded glomeruli to a given odorant class (see Methods). We excluded the mixed odorant class sensitivity vector from this analysis as it only contained one odorant, methyl anthranilate. Surprisingly, we found that the individual principal components of GNN Shapley fields are significantly correlated (see Methods) with only one (8 out of 10 first PCs) or two (one out of 10 first PCs) odorant class sensitivity vectors (*p <* 0.05) (Figure 4E). This is not an expected property of the Shapley fields, as GNN features were derived from a graph neural network trained on human perception, combined with molecular graph, yet the first ten principal components of the Shapley fields on the mouse OB often corresponded to a single “common chemical class” used in the set of stimuli. The same analyses applied to the PubChem Shapley fields proved less related to individual odorant chemical classes (Figure 4F), with 3 out of 10 first PCs correlated with just one chemical class. See Supplementary Table 1 for the actual correlation values. Note that the PubChem features show an approximate clustering of the odorant classes used in our experiment, whereas there is barely any clustering by odorant classes in the GNN feature space (SuppFig 8). Surprisingly, the GNN Shapley fields showed more pronounced one-to-one correspondence between principal components and chemical classes as compared to the PubChem Shapley fields, despite the lack of significant clustering of the GNN chemical features.

While there was no significant overperformance of the GNN-based prediction over the PubChem feature-based predictions (Figure 3), it seems that the principal components of the GNN Shapley fields are more consistent with the commonly used chemical classification. This raises the possibility that the human descriptors like “spicy” or “fruity” that were used in training the GNN model, together with the mouse OR responses and the molecular graphs used in the GNN, effectively yield the common classification of the odorant molecules (SuppFig 7). We speculate that this is because mouse olfactory perception is still sufficiently close to human olfactory perception despite the extensive evolutionary divergence in the olfactory systems of these two species.

## Discussion

Motivated by investigating odorant chemical feature tuning in the OB, we introduced an application of Shapley fields to explore the sensitivity of glomeruli to chemical features as a function of their position on the OB surface. The Shapley fields serve as an olfactory analogue of receptive fields in vision and other sensory areas. While chemical response spectra define receptive fields in the odorant space, they do not correspond to receptive fields in the chemical feature space, which Shapley fields aim to characterize as a more nuanced representation of glomerular sensitivity. Using two different computational approaches, we confirmed that the chemical receptive properties of glomeruli in the OB are consistent with a weak chemotopy in the organization of odorant representations across the OB surface. This agrees with earlier studies in Rubin and Katz (1999); Burton et al. (2022); Johnson et al. (1999); Johnson and Leon (2000); Johnson et al. (2005); Johnson and Leon (2007); Matsumoto et al. (2010); Meister and Bonhoeffer (2001); Soelter et al. (2020); Takahashi et al. (2004); Uchida et al. (2000) and at least partially addresses the concerns raised in later works Ma et al. (2012); Chae et al. (2019); Soucy et al. (2009).

To investigate how our conclusions are influenced by inherently different choices of the chemical feature sets, we repeated our analyses using an artificially constructed chemical feature set that combined molecular graph information with human perceptual data, using methods developed in Lee et al. (2023); Achebouche et al. (2022); Qian et al. (2023). We found that the outperformance of the Chemotopy vs. Chemotuning model does not depend on the choice of chemical feature sets, and that both chemical feature sets describe the receptive properties of the OB glomeruli with comparable accuracy. Note that unlike the findings in Qian et al. (2023), the predictions based on the GNN features were not significantly more accurate than those based on the PubChem features.

Furthermore, the Shapley fields derived from the two feature sets considered were strongly correlated, suggesting that the wide array of chemical features in PubChem and chemical features derived from human perception cover similar properties of mouse olfactory perception, as represented by the neural activity in the dorsal OB. This suggests that if one chooses a sufficiently diverse set of chemical features, the corresponding Shapley fields would capture the same receptive properties of the OB glomeruli and thus may provide a feature set-agnostic tool for investigating olfactory perception.

We also found that each principal component of the Shapley fields often significantly correlates with a single commonly accepted chemical class used in the presented stimuli. This suggests that these chemical classes could have been “discovered” based on the mouse olfactory responses.

## Materials and Methods

### Data collection

Data were collected from 2 male and 1 female compound heterozygous crosses of OMP-IRES-Cre (Jackson Laboratory stock #006668) and Rosa26-CAG-lox-STOP-lox-GCaMP6f (RCL-GCaMP6f; Ai95D; Jackson Laboratory stock #024105) mouse lines, aged 16 − 18 weeks. Prior to data collection mice were housed up to 5 per cage on a 12-hour light/dark cycle with food and water available ad libitum. Odorant responses were collected with epifluorescence imaging in isoflurane-anesthetized mice outfitted with a double tracheotomy to decouple respiration and anesthetic delivery from odorant presentation and sampling, using procedures identical to those described in Burton et al. (2019). All procedures were performed following the National Institutes of Health Guide for the Care and Use of Laboratory Animals and were approved by the University of Utah Institutional Animal Care and Use Committee.

### Olfactometry

Odorants were obtained from Sigma-Aldrich, TCI America, Bedoukian Research, or ICN Biomedicals and delivered using a custom olfactometer that allowed mixing of odorant vapor in free space prior and computer-controlled timing and choice of odorant identity Burton et al. (2019). Odorant delivery parameters were identical to those used in Burton et al. (2022) except that the end-stage eductor was omitted. Odorants were pre-diluted in caprylic/capric medium chain triglyceride oil (C3465, Spectrum Chemical Mfg. Corp.) as described in Burton et al. (2022) using dilutions ranging from 1 : 10 − 1 : 50 in order to achieve estimated delivered concentrations ranging from 1 − 1000 nM (median: 100 nM, interquartile range: 20 − 600 nM).

### Data processing

Odorant response matrices (odorant x glomerulus) were generated using the analysis pipeline described in Burton et al. (2022). Briefly, averaged Δ*F* response maps were generated from 4 − 6 repeated presentations of each odorant, after which regions of interest (ROIs) were selected manually using custom visualization software written in MATLAB (https://github.com/WachowiakLab). ROIs representing presumed individual glomeruli from the dorsal OB were segmented manually, with the segmenter blind to odorant identity Burton et al. (2022). Note that ROIs were drawn either following the rough contours of activation or with stereotyped-sized squares roughly centered within the contours of activation. After segmentation, response matrices were generated using the mean Δ*F* value across all pixels in an ROI, for all 59 tested odorants. Responses to 7 of the amine-class odorants were included *only* for two of the four OBs (1 mouse), due to concern about decomposition and the effective delivered concentrations of these compounds for the remaining two animals.

### Random Forests

The random forest binary classifier models were trained using the scikit-learn python library Pedregosa et al. (2011). The hyperparameters we were optimizing for were n_estimators (the number of trees in the forest), max_features (the number of features to consider when looking for the best split) and max_samples (the number of samples to draw from data to train each base estimator). Every other hyperparameter was chosen to be the default one given by the random forest classifier constructor, except the class_weight hyperparameter. Since we are dealing with heavily imbalanced datasets, the class_weight was set to “balanced_subsample”. Model performances were measured with the AUROC (Area Under Receiver Operating Characteristic curve). The cross-validation details are described in the relevant model descriptions.

### Chemotuning model

The Chemotuning model predicts glomerular activation from the chemical features of the odorants alone. More specifically, given a glomerulus-odorant pair (*g, o*), a random forest binary classifier predicts activation of *g* to *o* based on the chemical features of *o*. Thus, the Chemotuning model is a collection of random forest models predicting activation, one for each glomerulus on the OB. The hyperparameters for the RF that were optimized with respect to running time and performance were: n_estimators=128, max_features=“sqrt”, max_samples=0.5, class_weight=“balanced_subsample”.

### Chemotopy model

The Chemotopy model predicts glomerular activation from the chemical features of the odorants and position information about the glomeruli. More specifically, given a glomerulus-odorant pair (*g, o*), a random forest binary classifier predicts activation of *g* to *o* based on the chemical features of *o*, and the position of *g*. Thus, the Chemotopy model is a single random forest model for predicting activation of all the observed glomeruli. To compare with the Chemotuning model (see below) we added an additional feature, which is a random integer identity for each glomerulus. That is, we wanted to make sure the models are trained on the feature space of the same dimension and then compared. The prediction is cross validated across 10 folds and the AUROC metric is used to evaluate performance. The hyperparameters for the random forest that were optimized with respect to running time and performance were as follows: n_estimators=256, max_features=1.0, max_samples=0.7, class_weight=“balanced_subsample”.

### Peer prediction model

The peer prediction model predicts glomerular activation from the activation of its peers (the other glomeruli on the OB). More specifically, given a glomerulus-odorant pair (*g, o*), a random forest binary classifier predicts activation of *g* to *o* using as features the activations of all glomeruli except *g*, to *o*. Thus, the peer prediction model is a collection of random forest models predicting activation, one for each glomerulus on the OB. The model is cross validated in a leave-one-out fashion and the AUROC metric is used to measure performance. The hyperparameters for the random forest were optimized with respect to running time and performance and were the same as the ones for the Chemotuning model.

### Position of glomeruli

Each glomerulus was approximated by a minimal rectangle that encloses the region of interest (ROI) associated to the glomerulus. The center of the rectangle was taken to be the position of the glomerulus. For each OB, the *x* and *y* coordinates of the glomeruli were normalized to both have mean 0 and variance 1.

### The PubChem Substructure Fingerprint

The PubChem System generates a binary substructure fingerprint for chemical structures Kim et al. (2022). When considering unique and nonzero PubChem fingerprints for our datasets we ended up with 140 features for the 59 odorants dataset and 132 features for the 52 odorants dataset.

### PCA PubChem features

PCA dimensionality reduction was performed on the PubChem Substructure Fingerprint binary features with 59 and 52 odorants. Both were projected to 40 dimensions. The first 40 principal components accounted for around 90% of the variance in both datasets.

### GNN features

We used a dataset of 5955 small molecules that were made publicly available, after pre-processing, by the authors in Achebouche et al. (2022). Each molecule comes with a set of human perceptual odor labels. Originally these molecules, with their odor labels, were obtained from The Good Scents Company (TGSC) Database Company (2023), and Leffingwell Database Leffingwell and Associates (2012). The 160 odor labels were grouped into 21 categories and used for training a multitask binary classification Graph Neural Network (GNN) Scarselli et al. (2009); Zhou et al. (2020). We trained a multitask binary classification GNN using the DeepChem library Ramsundar et al. (2019) and the same network architecture as the one specified in Achebouche et al. (2022). Furthermore, we cross validated our GNN model across 5 folds and evaluated its performance with the AUPRC (Area Under Precision-Recall Curve) score. Our mean AUPRC score of approximately 0.40 agreed with the reported AUPRC score in Achebouche et al. (2022). We then trained a network by using the whole human perceptual dataset of 5955 odorants. Inspired by the idea of transfer learning Tan et al. (2018), we produced learned GNN features for our smaller dataset of odorants. That is, we used the learned weights in the network to human-derived learned features for our odorants, by considering their representations in the penultimate layer of the network, before the multitask softmax classification layer. We call these the GNN features. There were 256 GNN features in total. All the odorants in our datasets were present in the human perceptual dataset, except for 7; 6 of the 7 amines and methyl salicylate.

### Shapley values

Shapley values are a game theoretic approach to explain the outputs of machine learning models and we used the SHAP (SHapley Additive exPlanations) python library to compute them Lundberg and Lee (2017). Shapley values calculate the optimal credit allocation to features which locally explains the model outputs (Figure 1C). This is done by treating features as players in a cooperative game, and the model output as the game prize and then calculating the classic Shapley values from game theory Shapley (1951). For an arbitrary model, computing Shapley values has exponential time complexity and in practice Shapley values are only approximated. Instead, we used shap.TreeExplainer which uses dynamic programming to exactly compute Shapley values for tree-based models, in polynomial time Lundberg et al. (2020).

### Shapley rankings

Because Shapley values are associated with a given model, and the Chemotuning model is in fact a collection of individual models, one for each glomerulus, we ranked the chemical features using the Chemotopy model. Let 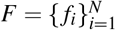 be any of the sets of chemical features that were described above. For a given an OB, we trained a Chemotopy random forest model, *M*, on every glomerulus-odorant pair (*g, o*) (no test/train split, no cross-validation) that predicted glomerular activation based on the position of *g* and chemical features of *o*. Let *f*_*i*_ denote a chemical feature in *F*, and *ϕ*_*i*_(*g, o*) denote the Shapley value of *f*_*i*_, which gives the contribution of *f*_*i*_(*o*) towards the prediction *M*(*g, o*). The global Shapley value of *f*_*i*_ is the mean absolute value of Shapley values of *f*_*i*_, over all observations (*g, o*), that is

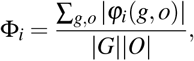

where |*G*| is the number of glomeruli on the OB, and |*O*| is the number of odorants. We thus have a collection 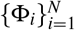 of nonnegative numbers that we rank in decreasing order. We call these ranks the Shapley rankings of the set of chemical features *F*. Note that we did not use a cross-validated training here, because we observed that the ranks didn’t change significantly, and the cross-validation procedure used for all the analyses in Figure 3 already insured that no overfitting had been possible.

Due to the randomness inherent to random forest models, a natural concern is the stability of these ranks. However, we found the Shapley ranks are for the most part stable with respect to the random choices of random forest models (SuppFig 5). The Shapley ranks of PubChem features were more stable than the ranks of GNN features. This is most likely due to GNN being a larger feature set (256 dimensions) as opposed to the PubChem features (140 or 132 dimensions) and hence there was more possibility of ranks changing during random forest training. The stability analysis was done by considering the Kendall rank correlation coefficients Kendall (1938) of Shapley rankings from different random seeds.

### Shapley fields

Given an OB, and a set of chemical features 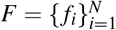, for each glomerulus *g* we train a Chemotuning model, *T*_*g*_, using the chemical features *F* (no train/test split, no cross-validation). Let *φ*_*i*_(*g, o*) be the Shapley value of *f*_*i*_, which gives the contribution of *f*_*i*_(*o*) towards the prediction 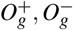 be the collections of odorants for which *φ*_*i*_(*g, o*) *≥* 0, *φ*_*i*_(*g, o*) *<* 0, respectively. Let

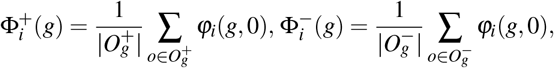

where 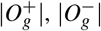 are the numbers of odorants in 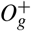 and 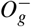, respectively. This construction yields, for each feature *f*_*i*_, two distributions on the OB which we call the positive and negative Shapley fields associated to *f*_*i*_. Intuitively, the Shapley fields represent the distribution of feature importance on the OB, very much analogous to receptive fields; one can also think of them as signed measures on the OB. We could have done a cross-validated training and made sure we add positive and negative Shapley values appropriately from different testing folds, and then define positive and negative Shapley fields. However, we observed that the values of the Shapley fields don’t change significantly and thus the additional computation time, which is not insignificant, would yield diminishing returns.

### Hierarchical clustering of Shapley fields

We concatenate the positive and negative Shapley fields, calculated from a fixed set of chemical features, from different OBs. Given the concatenated Shapley fields 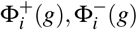, we perform a principal component dimensionality reduction and project to the first 20 principal components, which accounted for 70% (PubChem) or 50% (GNN) of the variance, see Figure 4D and SuppFig 3D. After the projection, we perform hierarchical ward linkage clustering with the cosine distance. The resulting dendrogram (Figure 2B and SuppFig 1B) and a height parameter are used to assign glomeruli to clusters.

### Odorant class sensitivity vectors

For each odorant class in our dataset, except “mixed”, (12 odorant classes in total) we define its odorant class sensitivity vector. It is a vector on the OB, meaning its coordinates are indexed by the glomeruli on the OB, and it counts to how many odorants from the given odorant class a given glomerulus was sensitive to. For example, if the odorant class in question is the acids, then the acid sensitivity vector counts for each glomerulus on the OB, to how many acids the glomerulus responded to. Since we used 5 acids in our experiments, this vector assigns values between 0 and 5 to each glomerulus.

### Correlations of Shapley fields principal components to odorant class sensitivity vectors

A principal component of Shapley fields from a chemical feature set is represented by a vector whose indices correspond to the glomeruli on the OB. This vector therefore lives in an *n*-dimensional space, where n is the number of glomeruli. The 12 odorant classes sensitivity vectors are also vectors in this *n*-dimensional space, and they span a 12-dimensional linear subspace. We measured the correlation between a principal component of Shapley fields and its projection to this linear subspace. The projection is a linear combination of odorant class tuning vectors with some weights. We calculated the statistical significance of these weights (*p <* 0.05) by randomly permutating the odorant identities that define the odorant class sensitivity vectors (*N* = 10000 trials). The statistically significant coefficients gave us the odorant classes’ tuning vectors the principal component of Shapley fields is correlated to.

### Distances between linear subspaces of Shapley fields

To compare Shapley fields induced by different chemical feature sets, we compared the linear subspaces spanned by *k* principal components they generate, where 1 ≤ *k ≤* 20. Here, each principal component is a high-dimensional vector whose coordinates are its values on each of the glomeruli (see Figure 4A). These *k*-dimensional subspaces can be represented as a matrix whose columns are the principal components. Given Shapley fields from different chemical feature sets we can then compare distances between these resulting linear subspaces, as we vary the PCA dimension k we are projecting to. The distances between these subspaces are calculated via principal angles, described below.

### Principal angles between subspaces

Angles between vectors in an *n*-dimensional space can be determined from the cosine-dot-product formula, i.e., if *θ* is the vector between angles *x* and *y* we have 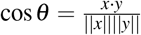. Treating vectors as the one-dimensional subspaces they span; this angle represents the angle between these one-dimensional subspaces. This idea extends to higher dimensions, and it is possible to define so-called principal angles between linear subspaces of any dimension. Specifically, given real-valued (*n × k*) matrices *A* and *B*, each one with pairwise orthogonal column vectors of length 1, one can find *k* principal angles *θ*_1_ ≤ *θ*_2_ ≤ … ≤ *θ*_*k*_ between the linear subspaces that are spanned by the column vectors of *A* and *B*. In practice the principal angles can be computed from the singular values of the matrix *A*^*T*^ *B* Björck and Golub (1973). *That is given σ*_*i*_, the *i*-th singular value of *A*^*T*^ *B*, we define cos *θ*_*i*_ = *σ*_*i*_.

### Chordal distance between subspaces

Given linear subspaces *V* and *W* of dimension *p*, let *θ*_1_ ≤*θ*_2_ ≤ … ≤ *θ*_*p*_ be the principal angles between *V* and *W*. The chordal distance between *V* and *W* is the quantity 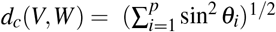, Conway et al. (1996). If we write *A* and *B* for matrices of column vectors of orthonormal basis vectors that generate *V* and *W*, respectively, by definition 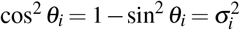, where *σ*_*i*_ is the *i*-th singular value of *C* = *A*^*T*^ *B*. In particular, 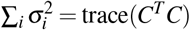. From this it follows that *d*_*c*_(*V*,*W*) = (*p−* trace(*C*^*T*^*C*))^1*/*2^.

### Spatial autocorrelation metrics

Let *N* be the number of spatial units indexed by *i* and *j*; *x* is the variable of interest; 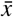 is the mean of 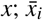 is the difference *x*_*i*_ *x*; *w*_*i* *j*_ is a matrix of spatial weights with zeroes on the diagonal; and *W* the sum of all *w*_*i*_ *j*. Then the Moran’s I and Geary’s C spatial autocorrelations are defined as: Moran’s I: Moran’s I is a measure of spatial autocorrelation that is given by the formula:

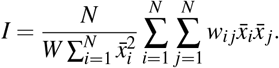

Geary’s C: Geary’s C is a measure of spatial autocorrelation that is given by the formula:

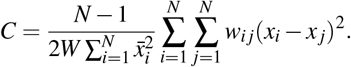

## Acknowledgments

The authors would like to thank Michael Schmuker for helpful discussions and suggestions. This work was supported by the NSF Next Generation Networks for Neuroscience Program, award 2014217, NIH grant F32MH115448, NSF grant 1555919, and NIH NINDS grant NS109979.

