## Supplemental figures for "Shapley Fields Reveal Chemotopic Organization in the Mouse Olfactory Bulb Across Diverse Chemical Feature Sets"

### Supplementary Materials

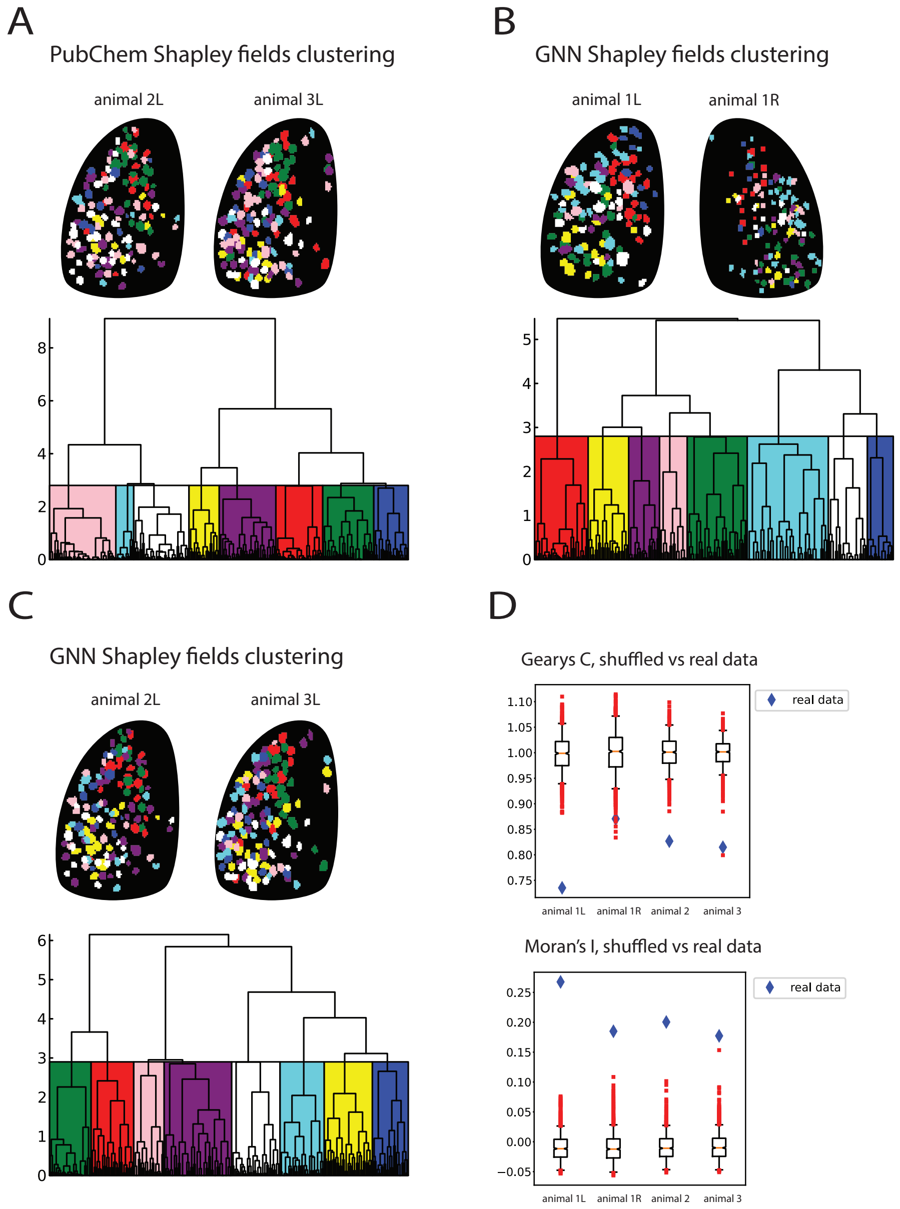


**SuppFig. 1: Mosaic chemotopic organization of feature importance on the OB. A. The clustering in the space of PubChem Shapley fields results in spatially correlated clusters on the surface of OB.** Hierarchical clustering of glomeruli on left OBs of animals 2 and 3, based on the first 20 principal components of Shapley fields, for the Pubchem chemical features. Cosine distance ward linkage clustering was used on the principal components of Shapley fields. A cutoff for clustering purposes was chosen as shown on the dendrogram. Most clusters have a "mosaic" distribution that is roughly consistent between the OBs. **B and C. The clustering in the space of GNN Shapley fields results in spatially correlated clusters on the surface of OB.** Hierarchical clustering of glomeruli on all OBs in our dataset based on the first 20 principal components of Shapley fields, for the GNN chemical features. Cosine distance ward linkage clustering was used on the principal components of Shapley fields. A cutoff for clustering purposes was chosen as shown on the dendrogram. Most clusters have a "mosaic" distribution that is roughly consistent between the OBs. **D. The spatial clustering inferred from GNN Shapley fields is statistically significant.** Glomeruli positions on the OB were randomly shuffled $n$ times ($n = 1000$). After each shuffling, a glomerulus was assigned to its original cluster from Panel B. The Moran's I and Gearys C clustering scores (Methods), across all 4 datasets show a highly significant difference ($p < 0.05$) between the clusters from the data (blue) and the randomly shuffled ones (red).

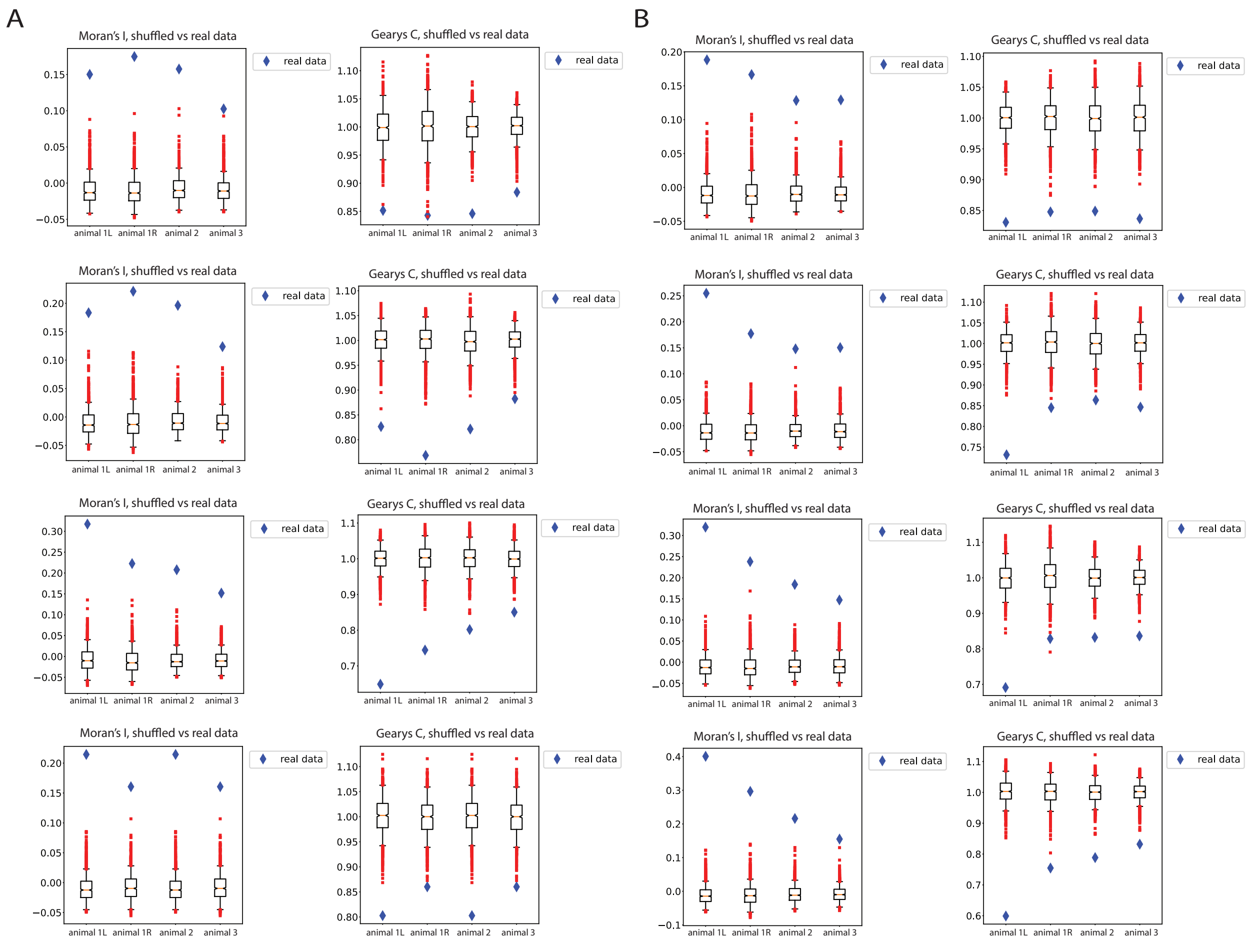


**SuppFig. 2: Statistical significance of clustering of Shapley fields. A. The spatial clustering inferred from PubChem Shapley fields is statistically significant.** Hierarchical clustering of glomeruli, based on the first 20 principal components of Shapley fields, for the Pubchem chemical features. Cosine distance ward linkage clustering was used on the principal components of Shapley fields. Different numbers of resulting clusters were considered (6,7,9,10 from top to bottom). Glomeruli positions on the OB were randomly shuffled $n$ times ($n = 1000$). After each shuffling, a glomerulus was assigned to its original cluster. The Moran's I and Gearys C clustering scores (Methods), across all 4 datasets show a highly significant difference ($p < 0.05$) between the clusters from the data (blue) and the randomly shuffled ones (red).**B. The spatial clustering inferred from GNN Shapley fields is statistically significant.** Same analysis as in Panel A, but for the GNN Shapley fields.

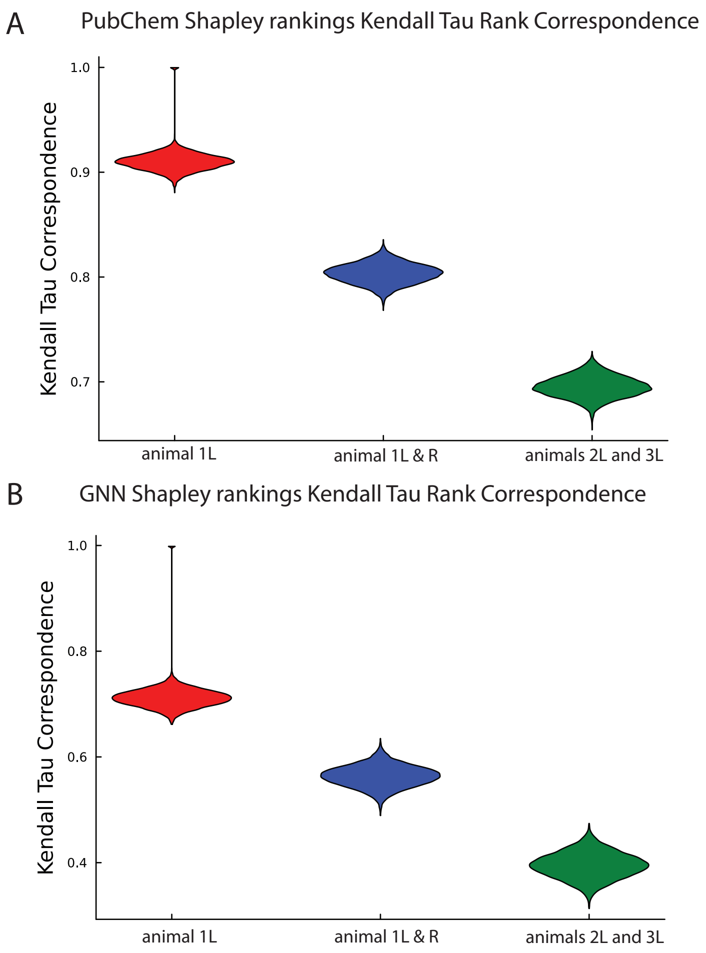


**SuppFig. 3: Stability of Shapley ranks of chemical feature sets. A. Kendall rank correlation coefficients of Shapley rankings of PubChem features.** Chemotopy model (see Methods) was trained 100 times (100 different random seeds), and the Shapley rankings of PubChem features used by the model were computed (see Methods). To analyze the stability of rankings (how much they change between random seeds) we computed the Kendall rank correlation coefficients between rankings. This comparison was done for Shapley rankings of PubChem features on the same OB (red), same animal but different OBs (blue), and on different OBs in different animals (green). Each violin plot contains 10000 such rank comparisons. The high values of Kendall rank correlation coefficients (the y-axis) indicate the stability of Shapley ranks of features. **B. Kendall rank correlation coefficients of Shapley rankings of GNN features.** Same analysis as in A, but for the the GNN feature set. Overall there is less stability for GNN Shapley ranks than for the PubChem Shapley ranks.

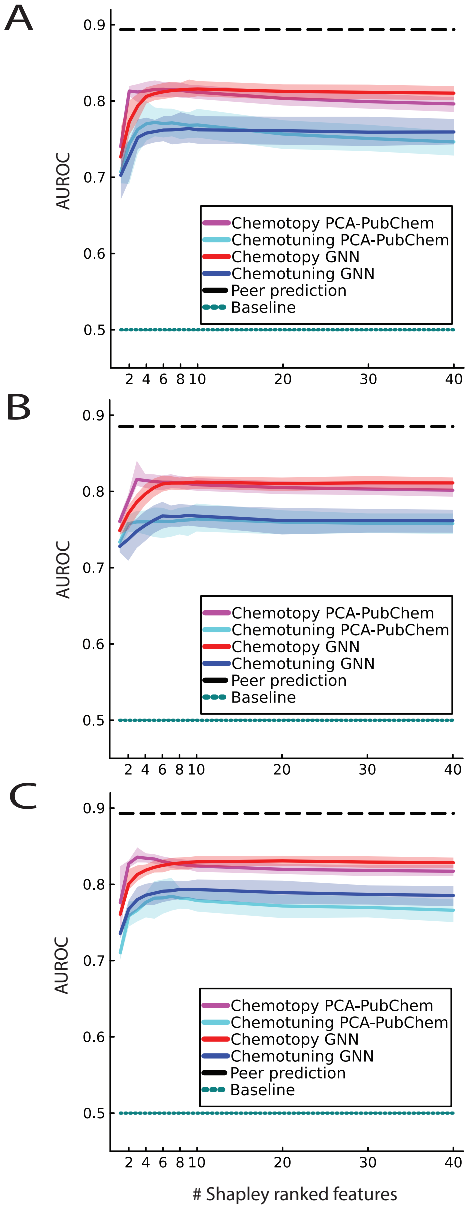


**SuppFig. 4: Chemotopy model outperforms chemotuning model. A. Three different models: chemotuning, chemotopy and the peer prediction are compared vs baseline, across chemical feature sets.** The features in a chemical feature set were ranked according to their global Shapley values, largest global Shapley value first (Methods). The features were added, one at a time according to their Shapley ranks, to the two models of glomerular activation prediction from chemical features; chemotopy and chemotuning (Methods). The mean cross-validated AUROC score is plotted along with its 95% confidence interval (shaded regions) against the number of Shapley ranked chemical features used. The mean cross-validated peer prediction model (black dashed line) AUROC score is also shown as well as the baseline (teal) having value of 0.5 which is the AUROC score of a random binary classifier. The 95% confidence interval for the peer prediction was too narrow to be perceptible on the plot ($0.8913 - 0.8962$). The data models performances shown are for the right OB of animal 1. **B. Three different models: chemotuning, chemotopy and the peer prediction are compared vs baseline, across chemical feature sets.** The same as in Panel A, but for the left OB of animal 2. **C. Three different models: chemotuning, chemotopy and the peer prediction are compared vs baseline, across chemical feature sets.** The same as in Panel A, but for the left OB of animal 3.

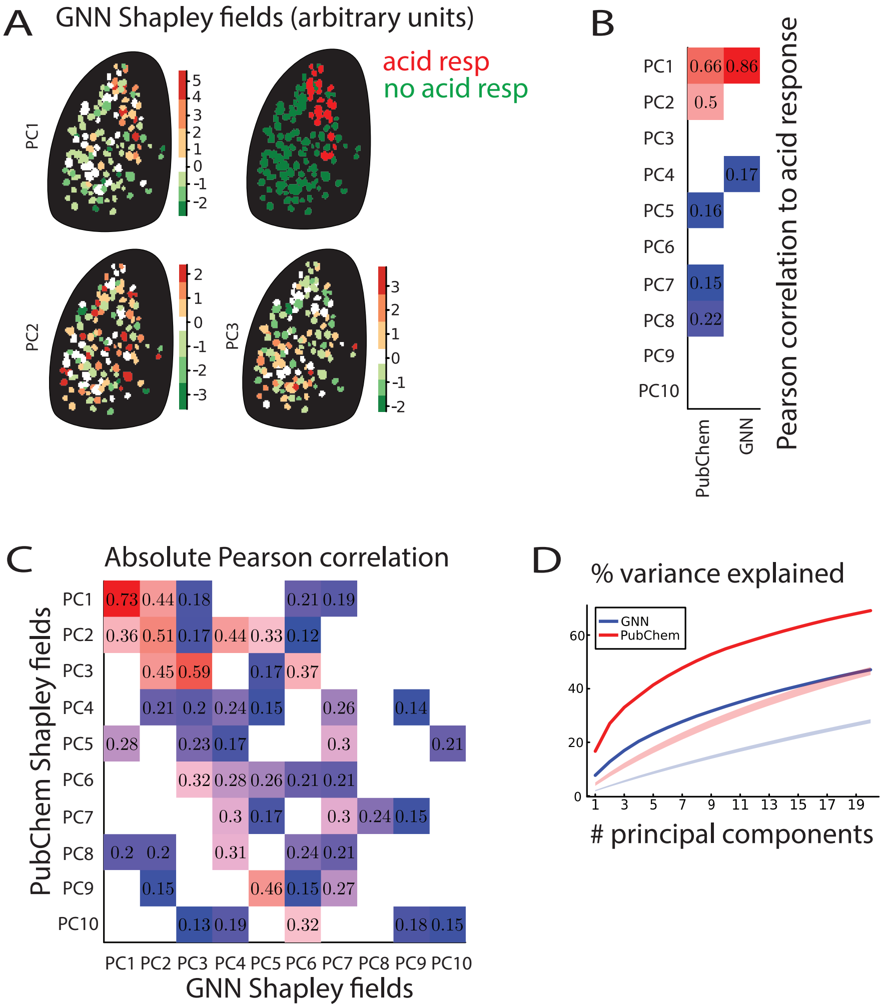


**SuppFig. 5: Shapley fields reveal an underlying "universal" chemical organization on the OB. A. Principal components of Shapley fields.** Importance of GNN features is projected on the OB via Shapley fields (Methods). First 3 principal components of Shapley fields on the left OB of animal 2, along with a classification of glomeruli into acid responsive (red) and acid non-responsive (green) are displayed. Visually the first principal component of GNN Shapley fields corresponds to acid tuning separation on the OB. **B. Interpretation of principal components of Shapley fields.** Absolute Pearson correlation values between first 10 principal components of PubChem and GNN Shapley fields with the responsiveness to acids vector. The responsiveness to acids vector is a count of the number of acids that activate a given glomerulus. Only significant correlations ($p < 0.05$) are shown. GNN Shapley fields PC1 is highly correlated with responsiveness to acids ($r = 0.86$). All the principal components of the GNN Shapley fields beyond PC2 show no significant correlation with acid response. In contrast, several PubChem Shapley fields principal components show correlation to responsiveness to acids to some degree, albeit none as high as the PC1 of GNN Shapley fields. **C. The Shapley fields of PubChem features and GNN features are highly correlated.** The absolute values of Pearson correlations of GNN's and PubChem's first 10 principal components of the respective Shapley fields are displayed. Only significant correlations ($p < 0.05$) are shown. There are significant degrees of correlation among the 10 principal components of Pubchem and GNN Shapley fields. **D. Variance explained by principal components.** We find that the first 20 principal components of PubChem Shapley fields account for about 69% of the total variance in Shapley fields (red curve). In contrast, the first 20 principal components of GNN Shapley fields account for about 47% (blue curve). The most probable reason for this difference is the simple difference in the number of Shapley fields; there are 264 PubChem Shapley fields and 512 GNN Shapley fields. The percentage of variance explained is statistically significant when compared to the shuffled Shapley fields principal components, from panel D, for both PubChem (red shaded region) and GNN (blue shaded region).

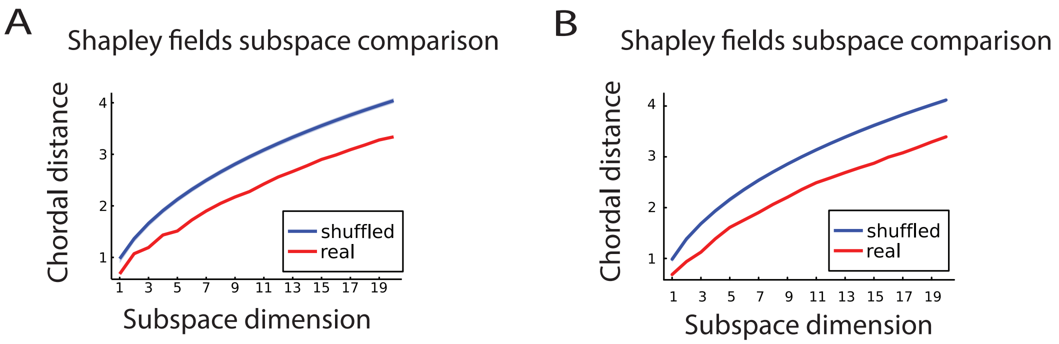


**SuppFig. 6: Human-derived chemical features and molecular fingerprints share chemical organization on the OB. A. Distances between linear subspaces of PCs of GNN and PubChem Shapley fields.** Principal angles and the resulting chordal distance between GNN and PubChem Shapley field principal component subspaces (Methods) were computed. We only considered the first 20 principal components of both GNN and PubChem Shapley fields and computed the principal angles of the linear subspaces they span, starting with PC1s and adding the other PCs one at a time. The resulting distance vs dimension curve is shown in red. We shuffled the activation patterns on both OBs of animal 1 ($N = 1000$), and recomputed both GNN and PubChem Shapley fields along with their chordal distances of subspaces generated by the first 20 principal components. The resulting curve (blue) with its 95% confidence interval shows that the real chordal distance between GNN and PubChem Shapley fields is statistically significant. **B. Distances between linear subspaces of PCs of GNN and PubChem Shapley fields.** Same analysis as in Panel A but for left OBs of animals 2 and 3.

| Odorants | acid | aldehyde | ester | furan | ketone | alcohol | amine | pyrazine | pyridine | phenolic | aromatic | thiazoline | mixed |
|---|---|---|---|---|---|---|---|---|---|---|---|---|---|
| Butyric acid | ■ | | | | | | | | | | | | |
| 2-Methylbutyric acid | ■ | | | | | | | | | | | | |
| Valeric acid | ■ | | | | | | | | | | | | |
| Isovaleric acid | ■ | | | | | | | | | | | | |
| Propionic acid | ■ | | | | | | | | | | | | |
| 2-Methylbutyraldehyde | | ■ | | | | | | | | | | | |
| 2-Methyl-2-pentenal | | ■ | | | | | | | | | | | |
| 2-Methyl-2-propenal | | ■ | | | | | | | | | | | |
| Butyraldehyde | | ■ | | | | | | | | | | | |
| trans-2-Methyl-2-butenal | | ■ | | | | | | | | | | | |
| Valeraldehyde | | ■ | | | | | | | | | | | |
| Butyl acetate | | | ■ | | | | | | | | | | |
| Methyl tiglate | | | ■ | | | | | | | | | | |
| Ethyl butyrate | | | ■ | | | | | | | | | | |
| Methyl valerate | | | ■ | | | | | | | | | | |
| Methyl 2-methylbutyrate | | | ■ | | | | | | | | | | |
| 5-ethyl-4-hydroxy-2-methyl-3(2H)-furanone | | | | ■ | | | | | | | | | |
| 5-methyl furfural | | | | ■ | | | | | | | | | |
| 2-Hexanone | | | | | ■ | | | | | | | | |
| 3-Octen-2-one | | | | | ■ | | | | | | | | |
| 3-Hepten-2-one | | | | | ■ | | | | | | | | |
| 2-Octanone | | | | | ■ | | | | | | | | |
| 4-Methyl-3-penten-2-one | | | | | ■ | | | | | | | | |
| 1-Hexanol | | | | | | ■ | | | | | | | |
| cis-3-Hexenol | | | | | | ■ | | | | | | | |
| Cyclohexylamine | | | | | | | ■ | | | | | | |
| 2-Methylbutylamine | | | | | | | ■ | | | | | | |
| N-Butyldimethylamine | | | | | | | ■ | | | | | | |
| N-Methyl piperidine | | | | | | | ■ | | | | | | |
| Butylamine | | | | | | | ■ | | | | | | |
| Octylamine | | | | | | | ■ | | | | | | |
| Cadaverine | | | | | | | ■ | | | | | | |
| 2,3-Diethylpyrazine | | | | | | | | ■ | | | | | |
| 2,3-Dimethylpyrazine | | | | | | | | ■ | | | | | |
| 5H-5-Methyl-6,7-dihydrocyclopenta[b]pyrazine | | | | | | | | ■ | | | | | |
| 2-Acetyl-3,(5 or 6)-dimethylpyrazine | | | | | | | | ■ | | | | | |
| 2-Methoxy-3-methylpyrazine | | | | | | | | ■ | | | | | |
| 2-Methoxy-3(5 or 6)-isopropylpyrazine | | | | | | | | ■ | | | | | |
| 2-Ethyl-3-methylpyrazine | | | | | | | | ■ | | | | | |
| 2-Isobutyl-3-methoxypyrazine | | | | | | | | ■ | | | | | |
| 2-Acetylpyridine | | | | | | | | | ■ | | | | |
| 4-tert-Butylpyridine | | | | | | | | | ■ | | | | |
| m-Cresol | | | | | | | | | | ■ | | | |
| Guaiacol | | | | | | | | | | ■ | | | |
| Acetophenone | | | | | | | | | | | ■ | | |
| 2-Hydroxyacetophenone | | | | | | | | | | | ■ | | |
| 4-Methoxyacetophenone | | | | | | | | | | | ■ | | |
| p-Anisaldehyde | | | | | | | | | | | ■ | | |
| Methyl salicylate | | | | | | | | | | | ■ | | |
| Propiophenone | | | | | | | | | | | ■ | | |
| Phenyl propionate | | | | | | | | | | | ■ | | |
| 2-Isobutylthiazole | | | | | | | | | | | | ■ | |
| Ethyl-2,5-dihydro-4-methylthiazole | | | | | | | | | | | | ■ | |
| 4-Methylthiazole | | | | | | | | | | | | ■ | |
| 2-Acetylthiazole | | | | | | | | | | | | ■ | |
| 2-Isopropyl-4-methylthiazole | | | | | | | | | | | | ■ | |
| 2-Methyl-2-thiazoline | | | | | | | | | | | | ■ | |
| 2,4,5-Trimethylthiazole | | | | | | | | | | | | ■ | |
| Methyl anthranilate | | | | | | | | | | | | | ■ |

Odorant classes

**SuppFig. 7: Odorant names and odorant classes used in experiments.**

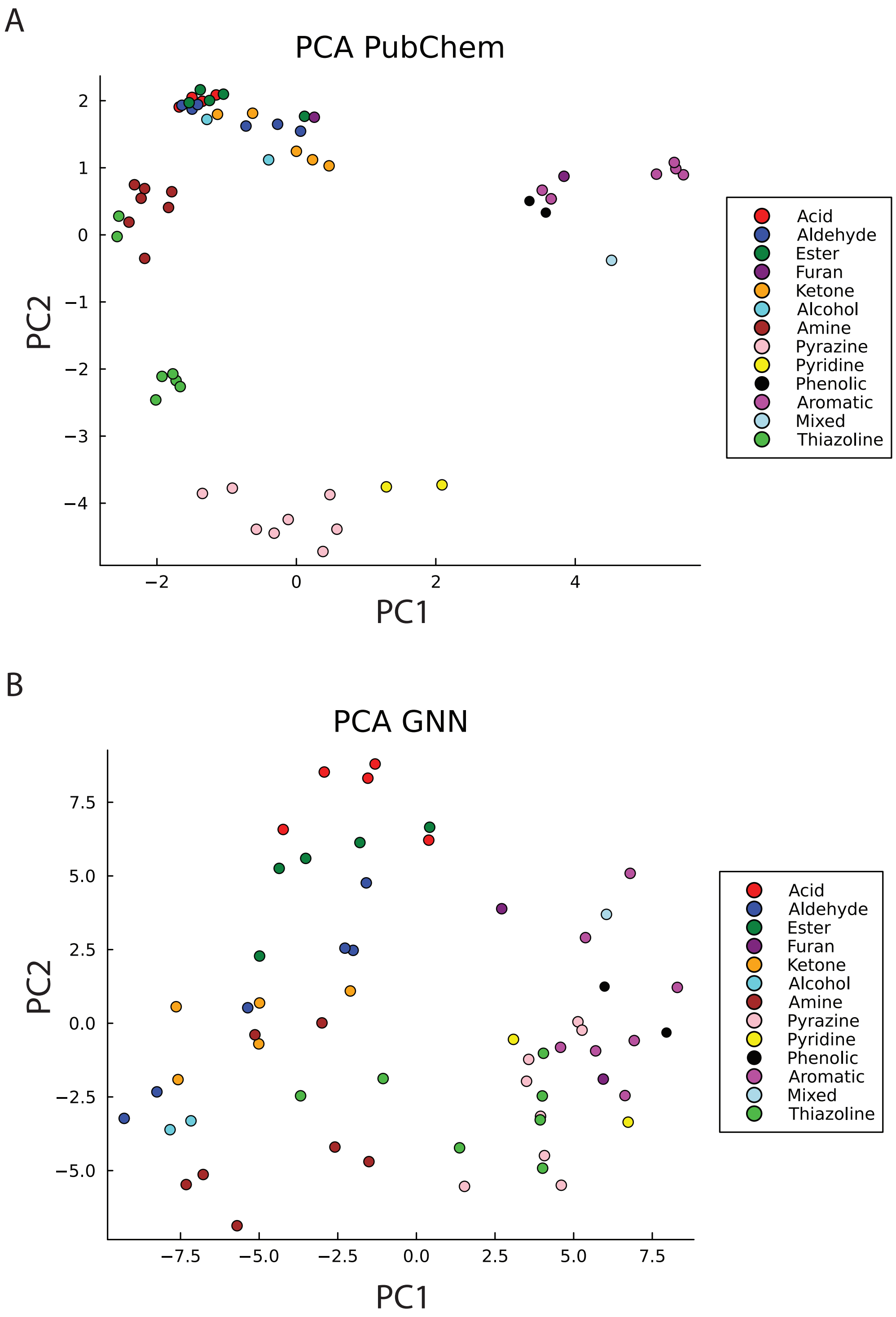


**SuppFig. 8: PCA projections of chemical feature sets. A. PCA projections of the PubChem feature set**. We reduced the dimension of the PubChem feature space of the 59 odorants used in our experiments to 2 dimensions, using PCA. Each odorant is represented as a point in the 2 dimensional space, with a color representing its odorant class. **B. PCA projections of the GNN feature set.** Same is in A, but for the GNN feature set.

| | PC1 | PC2 | PC3 | PC4 | PC5 | PC6 | PC7 | PC8 | PC9 | PC10 |
|---|---|---|---|---|---|---|---|---|---|---|
| PubChem Shapley fields | 0.87 p=4.2e-69 | 0.84 p=4.9e-61 | 0.81 p=4.2e-52 | 0.66 p=2.3e-29 | 0.87 p=2.6e-69 | 0.65 p=1.9e-28 | 0.5 p=1.6e-15 | 0.6 p=3.8e-23 | 0.51 p=3.5e-16 | 0.66 p=8.0e-29 |
| GNN Shapley fields | 0.93 p=1.0e-96 | 0.73 p=1.6e-38 | 0.78 p=2.9e-47 | 0.49 p=6.6e-15 | 0.73 p=6.6e-38 | 0.48 p=6.0e-14 | 0.47 p=1.9e-13 | 0.43 p=3.1e-11 | 0.5 p=1.7e-15 | 0.51 p=3.3e-16 |

**Supplementary Table 1: Correlations of Shapley fields to odorant classes sensitivity vectors.** Principal components of Shapley fields were projected onto the linear subspace spanned by odorant classes sensitivity vectors on the OB (Methods). The Pearson correlations between the principal components and their respective projections are shown in the table, along with the p-value of the correlation.
